# RECAP reveals the true statistical significance of ChIP-seq peak calls

**DOI:** 10.1101/260687

**Authors:** Justin G. Chitpin, Aseel Awdeh, Theodore J. Perkins

## Abstract

**Motivation:** ChIP-seq is used extensively to identify sites of transcription factor binding or regions of epigenetic modifications to the genome. A key step in ChIP-seq analysis is peak calling, where genomic regions enriched for ChIP versus control reads are identified. Many programs have been designed to solve this task, but nearly all fall into the statistical trap of using the data twice—once to determine candidate enriched regions, and again to assess enrichment by classical statistical hypothesis testing. This double use of the data invalidates the statistical significance assigned to enriched regions, and as a consequence, invalidates false discovery rate estimates. Thus, the true significance or reliability of peak calls remains unknown.

**Results:** Using simulated and real ChIP-seq data sets, we show that three well-known peak callers, MACS, SICER and diffReps, output optimistically biased p-values, and therefore optimistic false discovery rate estimates—in some cases, many orders of magnitude too optimistic. We propose a wrapper algorithm, RECAP, that uses resampling of ChIP-seq and control data to estimate and correct for biases built into peak calling algorithms. P-values recalibrated by RECAP are approximately uniformly distributed when applied to null hypothesis data, in which ChIP-seq and control come from the same genomic distributions. When applied to non-null data, RECAP p-values give a better estimate of the true statistical significance of candidate peaks and better false discovery rate estimates, which correlate better with empirical reproducibility. RECAP is a powerful new tool for assessing the true statistical significance of ChIP-seq peak calls.

**Availability:** The RECAP software is available on github at https://github.com/theodorejperkins/RECAP.

## 1 Introduction

Chromatin Immunopreciptation followed by high-throughput sequencing, or ChIP-seq, has become a central approach to mapping transcription factor-DNA binding sites and studying the epigenome [11, 21, 28]. ChIP-seq is the primary technique employed by a number of highly successful large-scale genomics projects, including ENCODE [5], modENCODE [25], the NIH Roadmap Epigenomics Project [17], and the International Human Epigenome Consortium [30]. Collectively, these projects have generated over 10,000 ChIP-seq data sets at a cost of 10s or 100s of millions of dollars, while other smaller-scale projects have generated many more. Many biological inferences are based on these data sets, including DNA binding motifs of transcription factors [16, 20], regulatory elements and networks [4, 12, 13], and possible connections to disease [27]. Thus, understanding exactly how much information we can or should extract from such data is a question of paramount importance.

Bioinformatics analysis of ChIP-seq data is a multi-stage process [18], with the end goal of identifying genomic regions of possible transcription factor-DNA binding, histone positions, chromatin marks, etc. There are numerous algorithms for identifying ChIP-seq enriched regions, or peak calling (e.g., [1, 7, 10, 24, 26, 29, 31, 32, 34, 36, 37]). Because ChIP-seq data is noisy, virtually all peak calling algorithms output peaks with associated p-values. These p-values are useful for ranking peaks in decreasing order of confidence, and estimating false discovery rates (FDRs) at different significance thresholds. But how well can we trust the p-values produced by peak callers?

For our study, we chose to focus on three peak callers: MACS (version 2.1.1.20160309) [8, 9, 37], SICER (version 1.1) [35, 36], and diffReps (version 1.55.6) [26]. We chose MACS because it is, at present, the most highly cited peak caller, and it is used by the ENCODE and modENCODE consortia for analysis of their data. SICER is another widely used and highly-cited algorithm, but one designed more for the detection of the broad, regional enrichment characteristic of certain chromatin marks. This suits some of our experiments below, although MACS is also able to detect such regions, particularly when used in “broad peak” mode. diffReps is designed to solve the differential enrichment problem—the comparison of two ChIP-seqs instead of a ChIP-seq and a control—which again comes up in some of our experiments.

Although these approaches to peak calling differ in a number of ways, all three (any many others from the list cited above) follow a common two-stage pattern: First, candidate peaks are identified by analyzing the ChIP-seq data, and second, those candidate peaks are evaluated for significance by comparing ChIP-seq data with some kind of control data. In the case of differential enriched region detection, two ChIP-seqs may be compared to each other by a similar process [26]. The problem with this design, as already pointed out by Lun and Smyth [19], is that it commits the statistical sin of using the data twice. The ChIP-seq data is used to construct hypotheses to test, the candidate peaks, and then the same ChIP-seq data, along with control or other ChIP-seq data, is used to test those hypotheses by means of classical statistical hypothesis testing. In general, when the hypothesis and the test both depend on the same data, classical p-values cannot be trusted.

When peaks’ p-values are wrong, it creates a host of other problems. For one thing, we no longer have a good basis for choosing a p-value cut off for reporting results. Relatedly, we do not know how much we can trust any given peak, or even the set of peaks as a whole. If a peak has a p-value of 10^−10^, we might feel that is very likely to indicate true transcription factor binding or epigenetic modification. But if the peak caller is biased, so that the real statistical significance of such a peak is only 10^−1^, then perhaps we should not put much stock in it after all. FDR estimates, which are also reported by most peak callers, are virtually meaningless when based on p-values that are themselves incorrect. Another problem arises if we try to compare results from different peak callers. To make comparisons “fair”, we might restrict both peak callers to the same raw p-value (or FDR) cut-off. But if one algorithm has highly biased p-values and the other does not, then this comparison will hardly be fair. Finally, any downstream analyses such as motif identification or regulatory network construction [4, 12, 13, 16, 20, 27] will be full of errors if we do not know which peaks are truly significant.

One approach to unbiased peak calling would be to develop a new peak calling approach from scratch, in a way that avoids double use of the data. However, as there are already many programs available that are satisfying in terms of identifying and ranking candidate peaks, with only their significance in question, we chose a different approach. We asked whether the p-values of peaks generated by these programs could be recalibrated to correct their bias. Happily, we found this to be largely possible through the new RECAP method that we introduce. RECAP stands both for the goal or our approach, <u>reca</u>librating <u>p</u>-values, and the method by which it is done, <u>re</u>sampling the read data and <u>ca</u>lling <u>p</u>eaks again. RECAP is a wrapper algorithm that is compatible with almost any peak caller, and in particular MACS, SICER and diffReps, for which we provide wrapping scripts. RECAP repeatedly resamples from ChIP-seq and control data according to a null hypothesis mixture. It then applies the peak caller to the resampled data, to sample the distribution of p-values under the null hypothesis of no difference between ChIP-seq and control. It uses the estimated cumulative distribution function of that distribution to adjust the p-values produced by the peak caller on the original data. We show that on a variety of different types of simulated null hypothesis ChIP-seq data, where there is no actual enrichment, RECAP-recalibrated p-values are approximately uniformly distributed between zero and one—as should be the case for well-calibrated statistical hypothesis testing. This gives a more intuitive way of choosing a significance cut-off for peak calling, and allows us to look at whether default cutoffs (such as the 10^−5^ raw p-value cutoff in MACS) are overly conservative or still too loose. FDR estimates based on recalibrated p-values are also more reliable, and in particular, we show that FDR q-values for peaks in ENCODE data track well the reproducibility of those peaks between biological replicates. In summary, RECAP allows for much more rigorous and rational analysis of the significance of enrichment in ChIP-seq data, while allowing researchers to continue using the peak calling algorithms they already prefer and have come to depend on.

## 2 Results

### 2.1 MACS, SICER and diffReps produce biased p-values

To test whether peak callers produce biased p-values, we generated 10 simulated null hypothesis data sets. In each data set, both ChIP-seq and control data comprise foreground regions and background regions. Foreground regions are 500bp long and spread approximately 20-25kb apart along a hg38-sized genome, and are the same for both ChIP-seq and control. Each ChIP-seq and control data set had 30,882,698 reads—one per 100 basepairs of the genome on average. 10% of the reads were placed uniformly randomly within the foreground regions, while the remainder were placed uniformly randomly within the background regions. Figure 1A shows a zoom-in on part of one of the randomly generated ChIP-seq data sets and its matching control. We chose these parameters for numbers of peaks, peak size and total reads to be broadly consistent with current ENCODE data sets.

**Figure 1:**
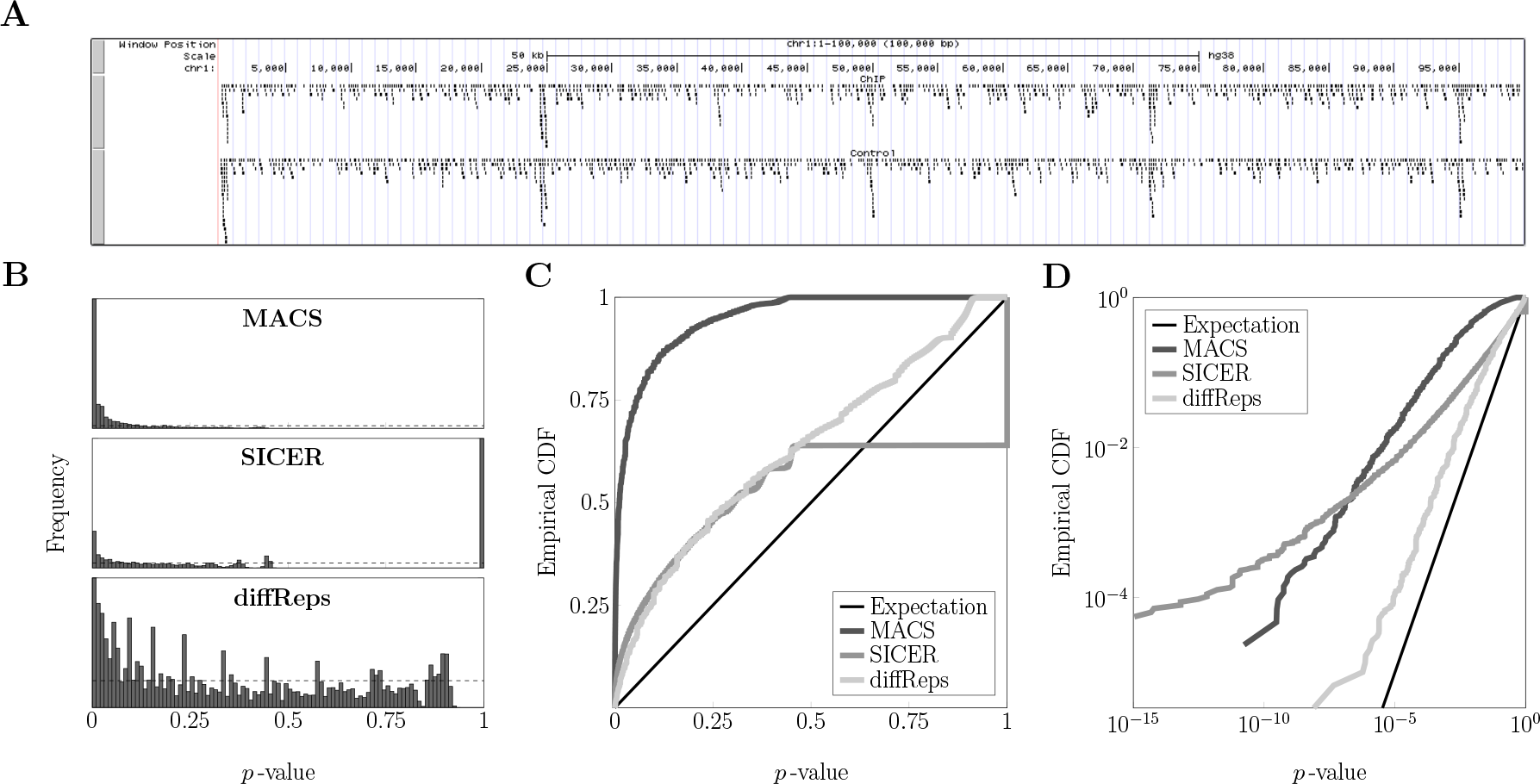
MACS, SICER and diffReps peak callers produce biased p-values. (A) Visualization of part of a simulated ChIP-seq read data set, with 500bp foreground regions every 20-25kbp, where read density is greater. Control data was generated similarly, with matching foreground regions, so a null hypothesis of no enrichment in ChIP-seq versus control is true for every possible genomic region.Peaks called by the three algorithms have p-values that are not uniformly distributed between zero and one, as should be the case for this null hypothesis data if p-values were well calibrated. (C,D) Empirical CDFs on linear (C) and log (D) axes also show the discrepancy from the uniform distribution.

We ran MACS, SICER and diffReps on these data sets, using default parameters with one exception. We set p-value or FDR cut-off thresholds at or as close as possible to 1.0, so that all candidate peaks, regardless of significance, would be reported. Figure 1B shows histograms of the p-values of the peaks produced by each program, for one of the 10 simulated ChIP-seq–control data set pairs. Results for the other 9 data sets were similar. By near universal definition, a p-value is the chance of observing data as or more “extreme” than some given data [33], under some null statistical model. As such, when applied to null-generated data, a well-calibrated method for calculating p-values should output p-values that are approximately uniformly distributed on [0,1]. That all three peak callers’ p-value distributions are non uniform is visually clear from Figure 1B, where the horizontal dashed lines indicate the uniform distribution, and from Figure 1C, where we plot the empirical CDFs of the p-values of the three programs. Well-calibrated p-values should have empirical CDF close to the thin black diagonal line. Although we will momentarily introduce a different statistic for quantifying deviation from uniformity, a simple KS-test shows that the three p-value distributions of the programs are statistically significantly different from the uniform distribution (*p* ≈ 0 incalculably small for all three).

Figure 1D shows the same empirical CDFs, but plotted on log-log axes. This plot is informative because most p-values are close to zero, and it is difficult to see their distribution on linear axes. Again, this plot shows that all three algorithms produce p-values that are non-uniformly distributed, and in particular, optimistically biased compared to the expectation under a uniform distribution of p-values. But it is now much more clear that diffReps’s p-values are the closest to being uniformly distributed, whereas MACS’s and SICER’s p-value distributions are farther afield. The curve for SICER, in fact, grows worse as p-value get smaller; SICER seems particularly prone to outputting highly significant p-values. Motivated by this log-log plot of empirical CDFs, we propose a measure of deviation from uniformity. For a given set of *N* p-values, we let *N*_1_/*N* be the fraction of those p-values in the range [0.1, 1], *N*_2_/*N* be the fraction in the range [0.01, 0.1), and more generally *N*_*i*_/*N* be the fraction in the range [10^−*i*^, 10^−*i*+1^). Then we quantify deviation from uniformity by the statistic: 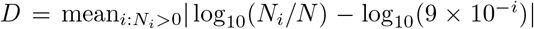. In words, this is the absolute difference between the logarithm of the fraction of peaks that should be in a p-value bin and the logarithm of the fraction of peaks that actually are in the bin, averaged over the non-empty bins. If a set of p-values is uniformly distributed on [0, 1], so that 90% of them fall in [0.1, 1], 9% fall in [0.01, 0.1), etc., then *D* evaluates to zero. Non-uniform distributions produce higher values of *D*. An advantage of this measure compared, for example, to the statistic used by the KS-test is that it pays equal attention to p-values at many different significance levels. In contrast, the KS-test looks at the maximum difference between the empirical CDF and the theoretical uniform CDF. For the SICER data, for example, this maximum occurs at *p* = 1, where approximately 40% of the peaks are. But the peaks with such high p-values are not of any biological interest, so it is undesirable for a performance metric to emphasize them to the exclusion of all else. For the present data, the deviations of the three algorithms’ p-value distributions evaluate to *D* ≈ 2.9 for MACS, *D* ≈ 4.1 for SICER, and *D* ≈ 0.8 for diffReps.

Although we will quantify bias and its removal more thoroughly in the next section, several important points remain regarding biases in the p-values produced by these programs. First, our results are not an artifact of the precise way the simulated null hypothesis ChIP-seq and control data sets were generated. For example, we also generated data with similar foreground regions but with 20% of reads in the foreground and 80% in the background. We also generated data with broad foreground regions of 4kbp containing 30% of the reads, leaving 70% for the background. For these data sets, we run MACS in broad peak mode. In all cases, we continue to see deviation from uniformity in the p-value distributions (Supplementary Figure 1A-C). Second, the amount of bias in these p-value distributions differs for the different types of data and for the different algorithms. This means that there is no universal correction that can be applied to the p-values, to bring them into line. Bias removal must operate in a way specific to the data being analyzed and to the program being used to call peaks.

Finally, it is important to note that evidence of bias can be seen in real data, not just simulated data. To show this, we turned to ChIP-seq data from the ENCODE consortium [5]. We chose to analyze data sets from the K562 cell line, as this is the cell line for which the most data sets are available. We identified all experiments that included two replicate ChIP-seq experiments and two matching controls (there were 88 such) and arbitrarily chose the first dozen of these for analysis (see Supplementary Table S1). In an attempt to approximate null hypothesis-like conditions, but using real data, we called peaks on each ChIP-seq data set using its ChIP-seq replicate as control. The resulting p-value CDFs for all three algorithms are shown in Supplementary Figure 1D-F. As with our simulated data, we see all the CDFs are optimistically biased, in some cases returning dramatic p-values exceeding 10^−300^. Thus, p-value bias is not just an artificial theoretical concern, but a genuine concern that is observable and should be expected in the analysis of real data.

### 2.2 RECAP: A wrapper algorithm that removes bias from peak caller p-values

Our approach to recalibrating p-values is based on empirically estimating an expected CDF for those p-values under a suitable null hypothesis. We put forth the null hypothesis that the ChIP-seq and control read data sets are drawn from the same distribution across the genome. That is, if we were to view each read as an i.i.d. sample where different positions on the genome would have different probabilities of being sampled, then we assume the sampling distribution of ChIP-seq and control are identical. Some work [3, 23] has explicitly attempted to estimate such distributions, but we will use a simpler mechanism for our p-value recalibration.

The RECAP algorithm is summarized below. From this point onward, we begin referring to the ChIP-seq data set as the “treatment” data. The reason for this is that the algorithm remixes ChIP-seq and control data into new data sets, and it would be confusing to call such remixed data sets by the name “ChIP-seq” when really they contain control data. (That said, we continue to call the control data by that name, as we know of no commonly-used term that could take its place.)

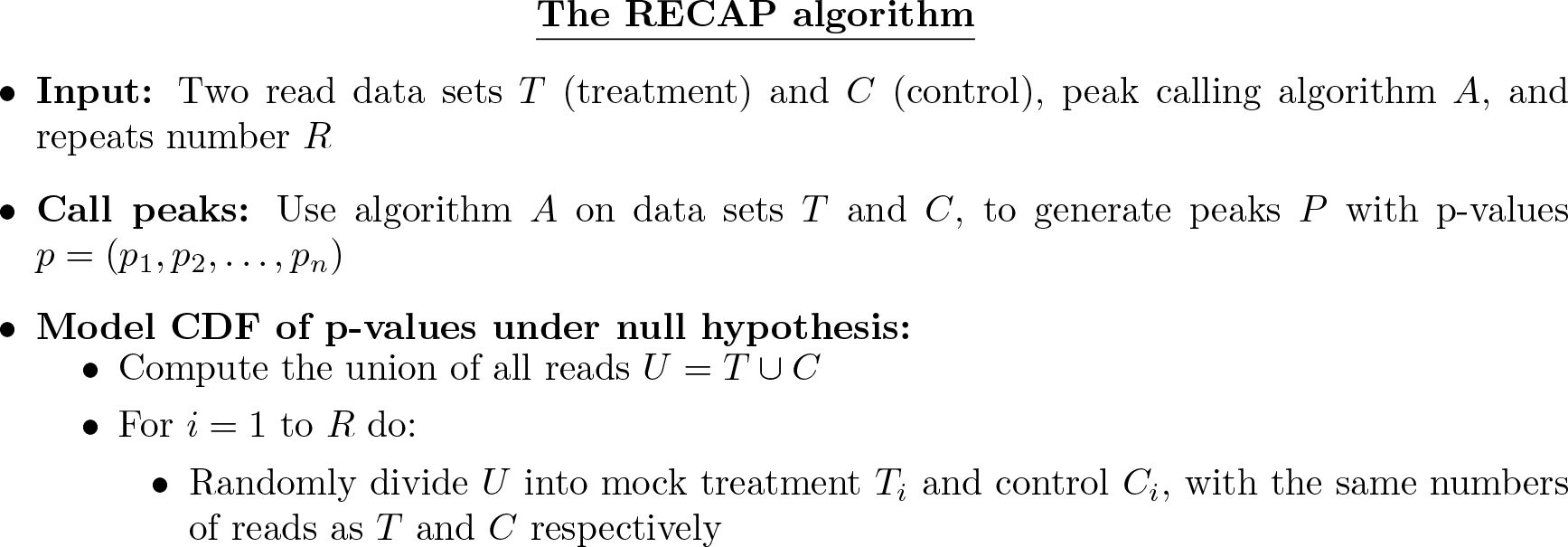

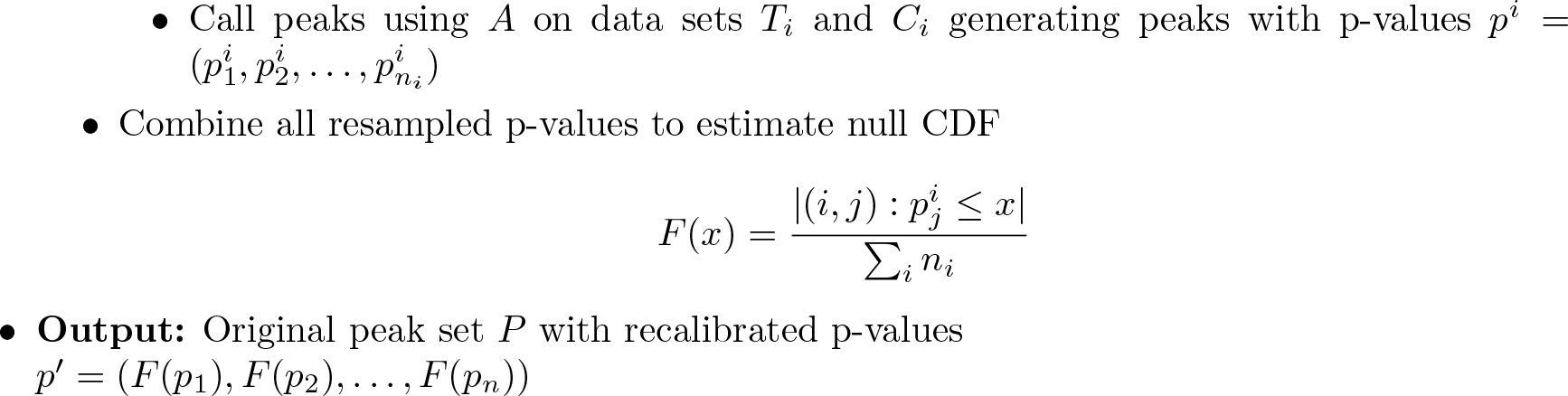

**Figure 2:**
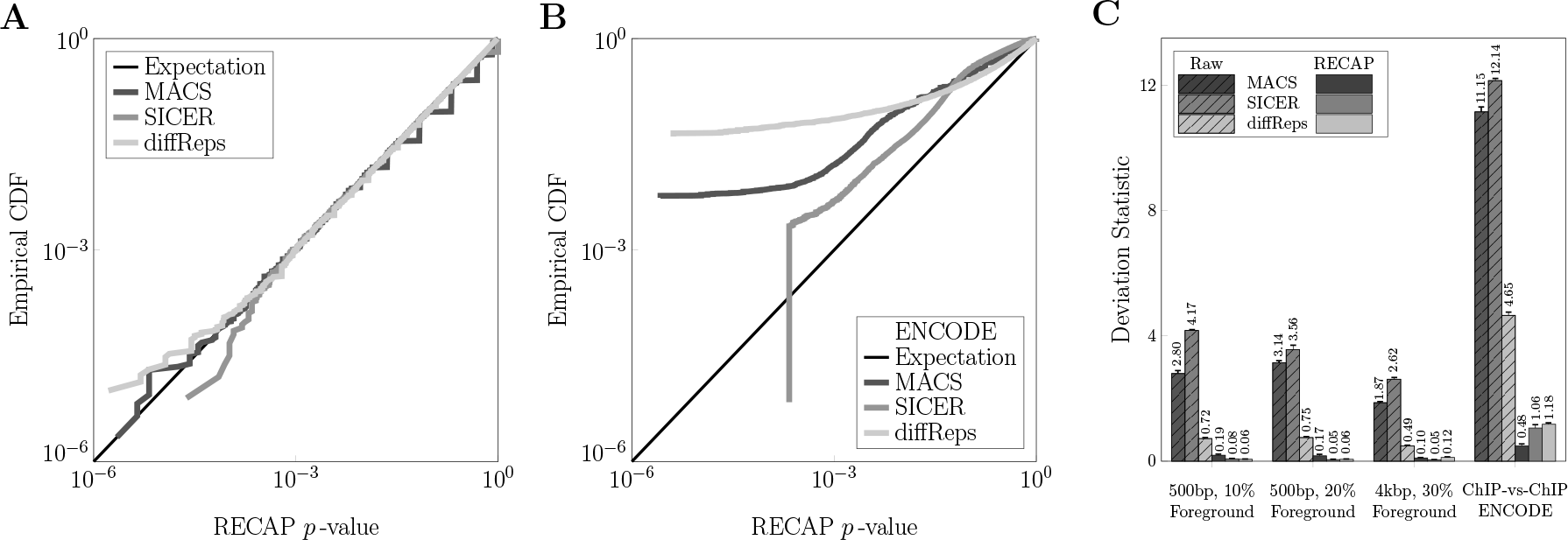
RECAP recalibrates peak callers’ p-values to a near-uniform distribution. (A) Log-log plot of the empirical CDF of recalibrated p-values for MACS, SICER and diffReps, on the simulated, 10% foreground, null-hypothesis data. (B) Reductions in deviation statistic, which measured difference from uniform distribution, for the RECAP recalibrated p-values for several types of simulated data (10 data sets each) and 12 matched pairs of ENCODE replicate ChIP-seq data.

The intuition behind the algorithm is that if the null hypothesis holds, we can simulate new-but-similar treatment and control data sets by resampling from the combined reads of the original treatment and control. If we do that repeatedly, and call peaks each time, we can estimate an average-case distribution of p-values for similarly-distributed data. Implicitly, this approach makes several assumptions. One assumption is that there even exists some notion of p-value distribution, given by *F*, that can be estimated. In principle, it is possible that every resampling of the data would generate peaks with radically different p-values or produce no peaks at all. If this were true, the “average” p-value distribution would not exist or would not be meaningful as a point of comparison for the original p-values *p*. In preliminary testing of all three algorithms, we found that while the *numbers* of peaks called could vary considerably between different resamples (particularly for MACS), the distributions of p-values were largely the same. Furthermore, a peak caller that did generate wildly different p-values for similar data sets would probably not be considered a good algorithm, due to lack of robustness. Second, our method assumes that every peak’s p-value in each of the *R* resamples can be viewed as i.i.d. samples from that distribution—justifying the standard empirical CDF estimate we use for *F*. In principle, because peak calling relies in part on local read densities, it is possible for nearby peaks to have non-statistically independent p-values. However, because these dependencies typically do not span a large portion of the genome, we expect the independence assumption is reasonable.

We tested RECAP’s ability to correct bias in peak p-values on a variety of simulated and real null hypothesis data sets. Figure 2A shows the results for the same 10%-reads, 500bp foreground region data set used for Figure 1B-D. Comparing particularly Figure 2A with Figure 1D, we see that RECAP has very substantially removed the bias. We obtained similar results when recalibrating p-values obtained from peak calling on replicate ENCODE ChIP-seq data sets; see Figure 2B for one of the real ChIP-seq pairs, and Supplementary Figure 2A-C for all 12 ChIP-seq pairs. Comparing to Supplementary Figure 1D-F, we see that bias has been very substantially reduced, although MACS and diffReps p-values remain somewhat optimistically biased. This could be the result of some genuine differences between the two replicates, causing peaks to appear in one and not the other, or causing overall signal fidelity to be different. For SICER, recalibration on some data sets results in optimistic p-values while others are pessimistic, although on average the curves cluster around the expectation line. A quantitative summary of bias before and after recalibration by RECAP is in Figure 2C. As was apparent visually, in all cases, we see that p-value distribution bias, as quantified by our deviation statistic *D*, is very substantially reduced.

### 2.3 Peak statistical signifance and FDRs estimated by RECAP

When p-values are not well-calibrated, the true statistical significance of individual peaks is unknown, and FDR estimates based on those p-values cannot be trusted. Conversely, if p-values are well-calibrated, then FDR estimates based on those p-values should also be well-calibrated. To examine these assertions, we ran further tests on both simulated and real (ENCODE) data. For the simulated data, we used as treatment the same data sets described above, but we used as control an equal number of reads distributed uniformly randomly across the genome. As such, there are many genuinely enriched regions in each simulated ChIP-seq data set. For the real data, we focused on the same 24 ENCODE ChIP-seq data sets mentioned in the previous section. We ran all three peak callers on each of the 24 independently, using matched controls as specified by the ENCODE project website (www.encodeproject.org; see Supplementary Table S2). We ran RECAP with either one or 10 resamplings to recalibrate the p-values. Based on the recalibrated p-values, we then computed q-values based on the method of Benjamini and Hochberg [2].

Figure 3A shows empirical CDFs of re-calibrated p-values for one typical simulated data set with 10% of reads in 500bp peaks. We observe that even after correction, the CDFs are significantly above the null expectation. This indicates genuine difference between treatment and control, which we know is correct for this data. Figure 3B shows similar behaviour for MACS on the ENCODE ChIP-seq data sets. (We focus on MACS results in this figure for the ENCODE data, because it is part of the ENCODE pipeline, and because we lack space to plot all three callers’ results. See Supplementary Figure 3A-B for the analogous plots for SICER and diffReps.) The p-value distributions shown in Figure 3B, where we call peaks on ChIP-seq versus control, are quite similar to the distributions shown in Supplementary Figure 2A, where we call peaks on ChIP-seq versus replicate ChIP-seq. However, there is a quantitative difference, as we summarize in Figure 3C. For all types of data, recalibration lowers the deviation statistics. However, it does not lower them as much as when we call simulated or real ChIP-seqs against each other (compare to Figure 2 and Supplementary Figure 2). This indicates, as one would expect, that although replicate ChIP-seqs may have some differences, there are more differences between ChIP-seqs and controls, and more or more strongly genuinely enriched regions.

**Figure 3:**
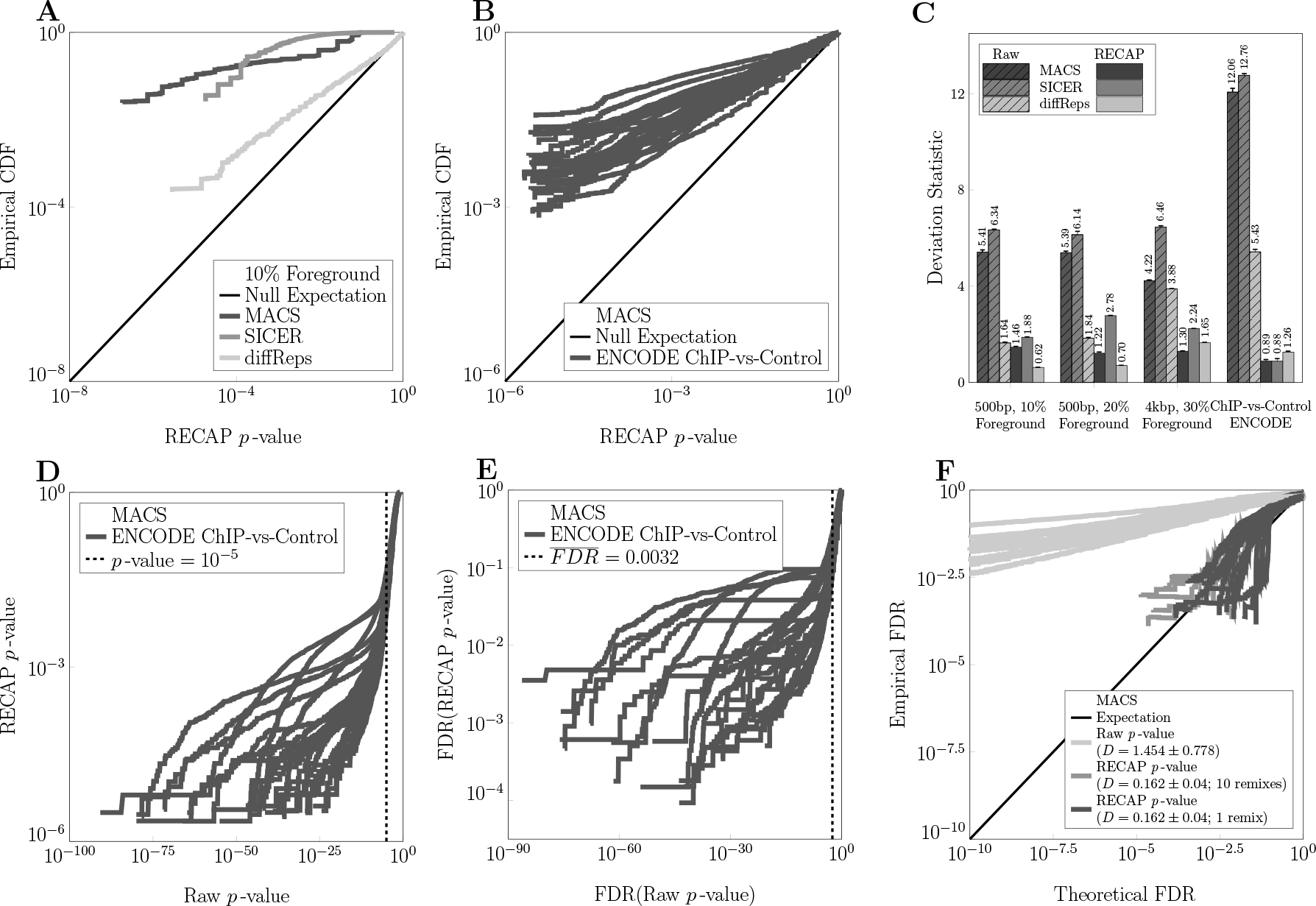
False discovery rate and reproducibility analyses based on recalibrated p-values. (A) Empirical CDF of recalibrated p-values for each algorithm on simulated non-null data. (B) Empirical CDFs of MACS’s p-values on ENCODE data sets. (C) Deviation statistics before and after recalibration. (D) Raw versus recalibrated p-values for MACS on ENCODE data. (E) FDR estimates based on recalibrated versus raw p-values for MACS on ENCODE data. (F) Peak reproducibility rates versus FDR estimates based on raw or recalibrated p-values.

Beyond the deviation statistics, a point of central interest is how p-values are transformed by calibration when there are genuinely enriched regions. Figure 3D and Supplementary Figure 3C-D plot recalibrated against raw p-values for MACS, SICER and diffReps. For MACS and SICER, peaks with phenomenal p-values like 10^−50^ have significance upon recalibration on the order of 10^−3^ to 10^−6^. While many of these may still be significant, their level of significance is not nearly what one might have expected. For diffReps, we find that the p-values, while optimistic, are not nearly so biased on the ENCODE data, and are typically recalibrated by an order of magnitude or less. For MACS, the default raw p-value cutoff is 10^−5^. Across the 24 ENCODE ChIP-seqs, we found that the least significant peak (i.e. the one with largest p-value less than or equal to 10^−5^) had a recalibrated p-value consistently near 10^−2^ (mean 0.0195, standard deviation 0.0077). If we apply the same raw p-value cutoff to SICER peaks, the least significant peaks have a recalibrated significance of 0.1065 ± 0.1807. For diffReps, however, these least significant peaks have fairly-correct raw p-values of 1.071 × 10^−4^ ± 1.717 × 10^−4^.

Figure 3E and Supplementary Figure 3E-F show FDR estimates (q-values) based on recalibrated versus uncorrected p-values. Again, we see a great discord between the two, particular for MACS and SICER. For MACS for example, uncorrected FDR estimates between 10^−30^ and 10^−60^—which would suggest no false positives at all in a typical set of peak calls—map to recalibrated FDR estimates between 10^−1^ and 10^−4^—suggesting a much higher level of false positives. If we look at the peaks up to an uncalibrated q-value threshold of 10^−5^, FDRs estimated based on our recalibrated p-values are 0.0032±0.0022 for MACS, 0.1583±0.1833 for SICER, and 1.2448 × 10^−4^±2.0310 × 10^−4^ for diffReps. Given that data sets such as these ENCODE ones typically have thousands or tens of thousands of peaks, the difference in uncalibrated and calibrated FDR estimates means the difference between essentially zero estimated false positive peaks and hundreds or even thousands of estimated false positive peaks.

Finally, we turn to the question of whether theoretical FDR estimates correlate to empirical FDRs. To do this for the ENCODE ChIP-seq replicate pairs, we designated a peak called in a ChIP-seq data set as a true positive if it overlaps (by as little as one basepair) a peak called in the replicate data set. Otherwise, a peak is designated a false positive. In Figure 3F and Supplementary Figure 3G-H, we plot empirical FDRs against FDR estimates based on either uncalibrated or recalibrated p-values. For MACS, theoretical FDRs based on uncorrected p-values are optimistic. For example, a theoretical FDR of 10^−5^, at which level there should be essentially no false positives, corresponds to an empirical FDR (i.e. reproducibility failure rate) of approximately 10^−2^, suggesting hundreds of peaks would be false positives. However, we see that theoretical FDRs based on recalibrated p-values track fairly well the empirical FDR, as seen by the clustering of the curves around the “expectation” line. Thus, for MACS, we suggest that FDR calculations based on recalibrated p-values can be trusted as a rough approximation of reproducibility, whereas FDRs based on uncalibrated p-values should not be taken at face value. For SICER the story is slightly more complicated. In Supplementary Figure 3G we see (and the deviation statistic confirms it) that FDR estimates based on recalibrated p-values more accurately reflect empiricial reproducibility. However, there is high variability across data sets, so that it is difficult to trust the results on any given data set. For diffReps (Supplementary Figure 3H), the bias in p-values was already low. Nevertheless, we see a modest improvement in the accuracy of FDR estimates based on recalibrated p-values versus estimates based on the raw p-values.

## Discussion

In this paper, we have looked at the question of how statistically significant peak calls in ChIP-seq data are. We argued that, for various reasons, peak callers have optimistic biases built into them, such that the actual significance of the called peaks is not clear. We simulated null hypothesis data with different amounts of background noise and with either narrow or broad foreground regions— regions where read densities are higher than in the rest of the genome, but equivalently high in treament and control, so that there is no differential enrichment. We documented optimistic bias in three widely-used peak callers, MACS, SICER and diffReps. Also importantly, we showed that the amount of bias differs between algorithms and between data sets, so that there is no simple universal correction that can be applied to correct the problem. With such miscalibration of p-values, we have no real accurate knowledge of the statistical significance of any given peak and, although this was not a focus of our paper, no way of comparing the significance of results from different approaches.

We then described RECAP, a wrapper algorithm that resamples from the combined treatment and control data to estimate p-value distributions when a null hypothesis of no differential enrichment is true. RECAP uses that information to compute a peak caller-specific and data set-specific correction to peak p-values when peaks are called on the treatment versus the control. We showed that RECAP can virtually eliminate bias in p-values generated from simulated null hypothesis data, and can largely eliminate bias in p-values from a collection of real ENCODE ChIP-seq data sets. We found that recalibrated p-values for all three algorithms are not nearly as statistically significant as the reported raw p-values—in some cases many orders of magnitude less significant—although it remains clear that there are many genuinely enriched regions. FDR estimates based on recalibrated p-values suggest that, with default peak calling parameters, FDRs may be approximately 100 times higher than previously estimated, although this would still correspond to a minority of the peaks being false positives in the ENCODE data sets we studied.

Although RECAP is a complete system as it stands, there are a number of possible directions for further work. First, while we showed our recalibrated p-values are useful for more accurate FDR estimation, they should also be useful for local FDR estimation [6]. While global FDR analysis tells us how many false positives may be in a given set of returned results, local FDR analysis can tell us how likely any individual peak is to be a false positive. Local FDR analysis requires a scheme for estimating null and non-null p-value distributions, and the prior probability of true versus false peaks. Second, we note that our p-value recalibration scheme monotonically transforms raw p-values based on an empirical CDF. This means that if one peak is more significant than another by raw p-value, its recalibrated p-value will also be more significant. But plausibly, some kinds of peaks or some genomic regions are more likely to be false positives than others. Indeed, other ongoing work in ChIP-seq analysis aims at uncovering and removing local biases in ChIP-seq signals that can unduly influence peak calling [15, 14, 22]. This suggests that peak-specific p-value corrections might be desirable, although it is unclear how this can best be done. Finally, although we have focused here on ChIP-seq peak calling, it is entirely reasonable to think that similar problems with p-value calibration may occur in other areas of high-throughput data analysis. For example, this may occur in DNA variant-calling, where complex conditions of uni- or bi-directional read coverage or other types of pre-filtering are sometimes applied before candidate variants are tested statistically. This double-usage of the data, to both select hypotheses for testing and to compute significance for those hypotheses, is a recipe for biased p-values. Perhaps in such cases, a similar read-resampling scheme could be used to calibrate p-values output by different variant callers.

## Supplementary information for

**Table S1:**
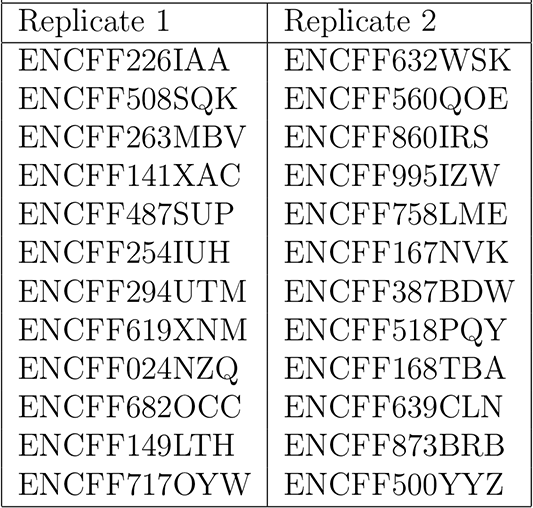
ENCODE identifiers for ChIP-seq replicate datasets

| Table S1: ENCODE identifiers for ChIP-seq replicate datasets |  |
| --- | --- |
| Replicate 1 | Replicate 2 |
| ENCFF226IAA | ENCFF632WSK |
| ENCFF508SQK | ENCFF560QOE |
| ENCFF263MBV | ENCFF860IRS |
| ENCFF141XAC | ENCFF995IZW |
| ENCFF487SUP | ENCFF758LME |
| ENCFF254IUH | ENCFF167NVK |
| ENCFF294UTM | ENCFF387BDW |
| ENCFF619XNM | ENCFF518PQY |
| ENCFF024NZQ | ENCFF168TBA |
| ENCFF682OCC | ENCFF639CLN |
| ENCFF149LTH | ENCFF873BRB |
| ENCFF717OYW | ENCFF500YYZ |

**Table S2:** ENCODE identifiers for ChIP-seq datasets and their controls

| Table S2: ENCODE identifiers for ChIP-seq datasets and their controls |  |
| --- | --- |
| ChIP-seq | Control |
| ENCFF226IAA | ENCFF910IKB |
| ENCFF632WSK | ENCFF227IZS |
| ENCFF508SQK | ENCFF910IKB |
| ENCFF560QOE | ENCFF227IZS |
| ENCFF263MBV | ENCFF227IZS |
| ENCFF860IRS | ENCFF910IKB |
| ENCFF141XAC | ENCFF227IZS |
| ENCFF995IZW | ENCFF910IKB |
| ENCFF487SUP | ENCFF910IKB |
| ENCFF758LME | ENCFF227IZS |
| ENCFF254IUH | ENCFF227IZS |
| ENCFF167NVK | ENCFF910IKB |
| ENCFF294UTM | ENCFF227IZS |
| ENCFF387BDW | ENCFF910IKB |
| ENCFF619XNM | ENCFF910IKB |
| ENCFF518PQY | ENCFF227IZS |
| ENCFF024NZQ | ENCFF910IKB |
| ENCFF168TBA | ENCFF227IZS |
| ENCFF682OCC | ENCFF227IZS |
| ENCFF639CLN | ENCFF910IKB |
| ENCFF149LTH | ENCFF910IKB |
| ENCFF873BRB | ENCFF227IZS |
| ENCFF717OYW | ENCFF910IKB |
| ENCFF500YYZ | ENCFF227IZS |

**Figure S1:**
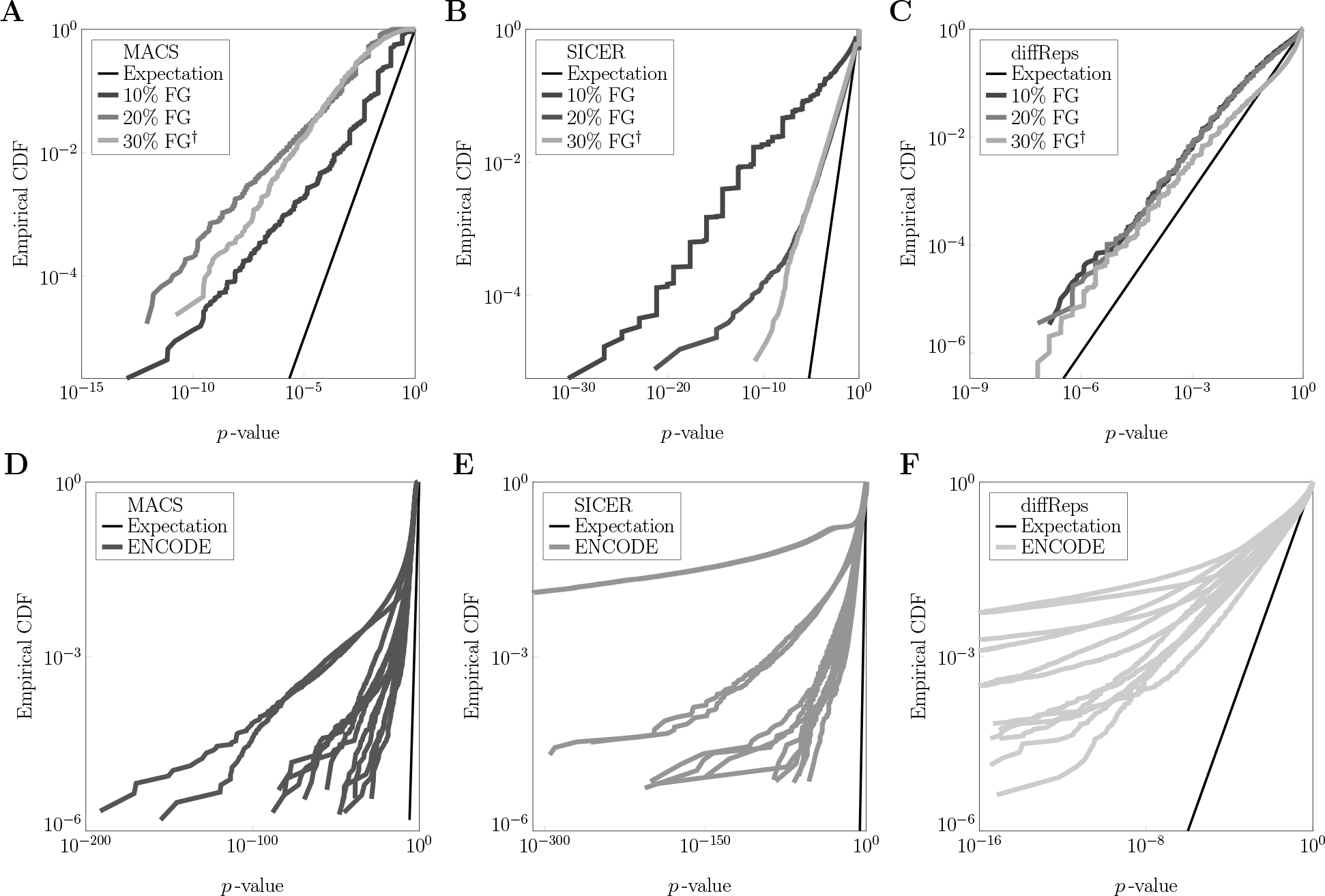
Empirical CDFs for peak raw p-values for MACS (A), SICER (B), and diffReps when applied to different types of simulated null hypothesis data: 500bp peaks with 10% of total reads, 500bp peaks with 20% of total reads, and 4kbp peaks with 30% of total reads. Empirical CDFs for peak raw p-values for MACS (D), SICER (E), and diffReps (F) when applied to ENCODE data, where one ChIP-seq replicate is used as control for another ChIP-seq replicate.

**Figure S2:**
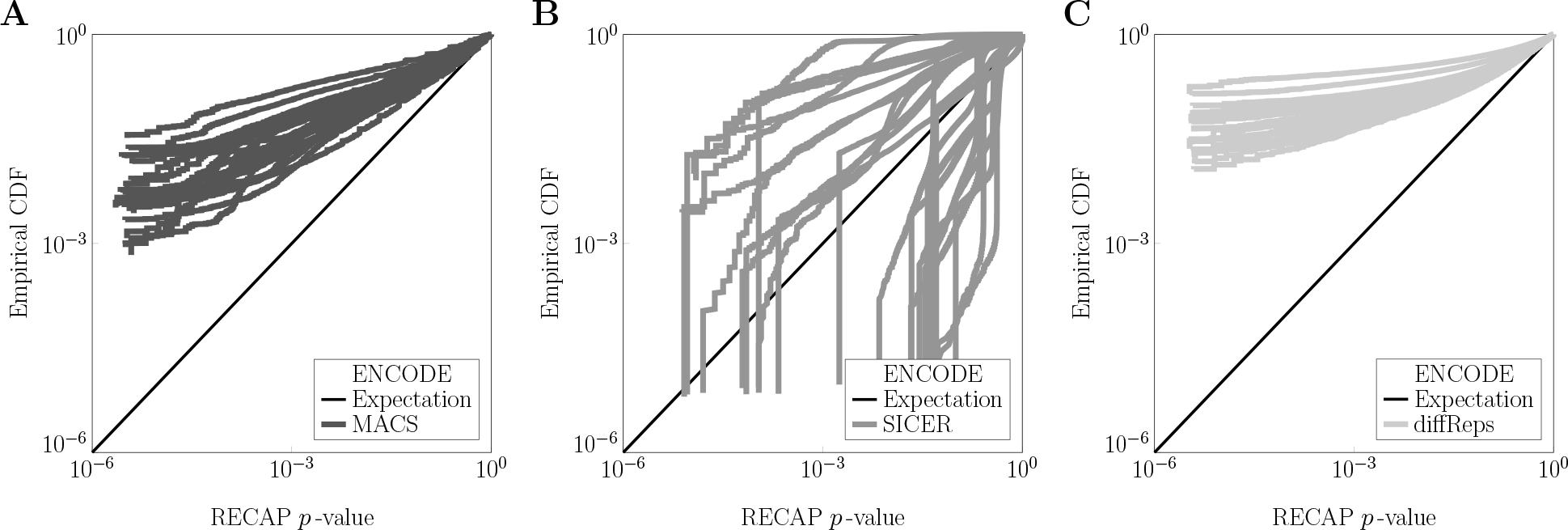
Empirical CDFs for peak recalibrated p-values for MACS (A), SICER (B) and diffReps (C) when applied to 12 replicate pairs of ENCODE ChIP-seq data.

**Figure S3:**
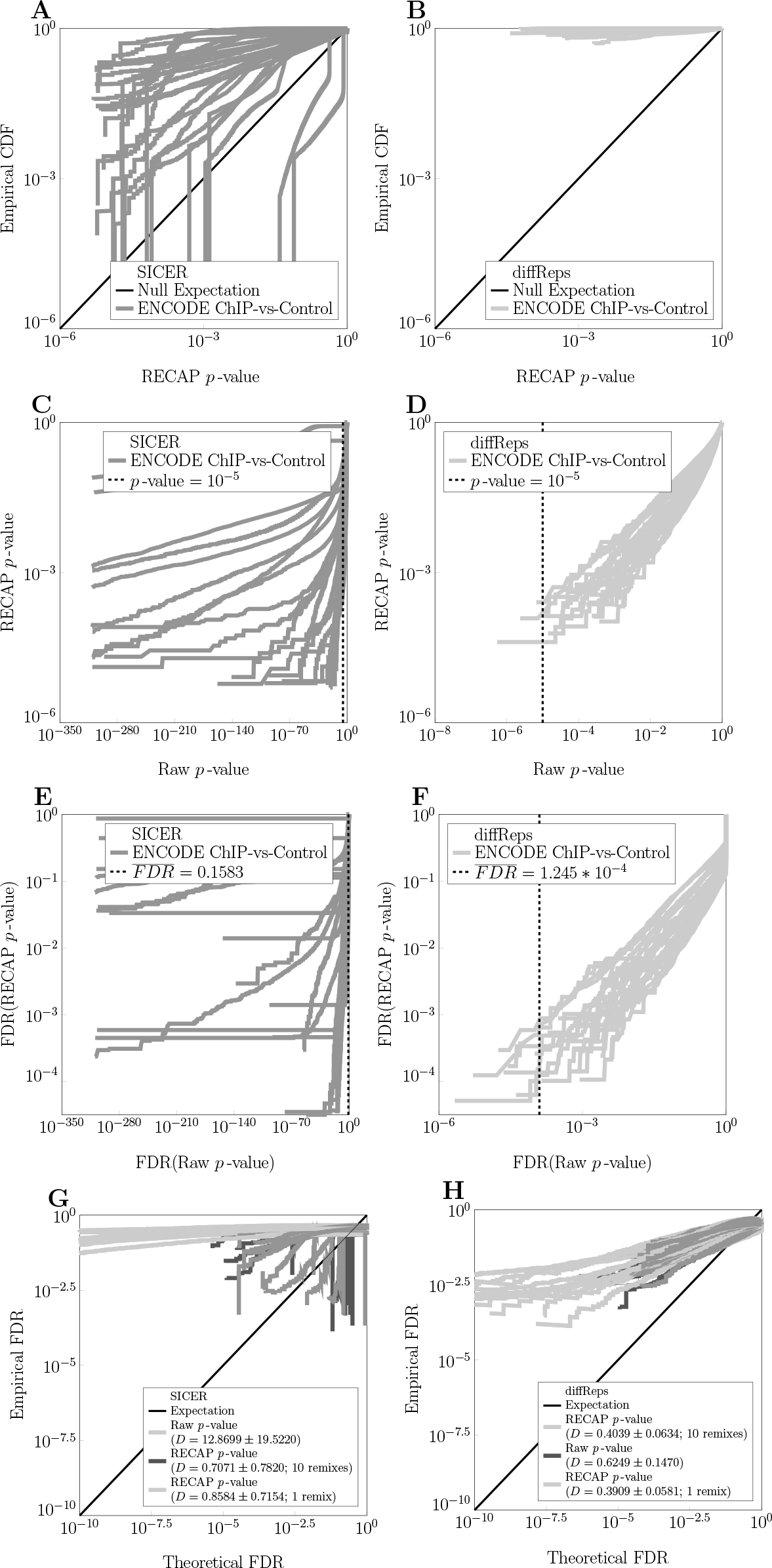
P-values and theoretical and empirical FDRs for SICER and diffReps for ENCODE ChIP-seq versus control peak calling.

